# Measles virus co-opts epithelial-to-mesenchymal transition and live cell extrusion to exit human airway epithelia

**DOI:** 10.1101/2024.07.12.603350

**Authors:** Camilla E. Hippee, Lorellin A. Durnell, Justin W. Kaufman, Eileen Murray, Brajesh K. Singh, Patrick L. Sinn

## Abstract

Measles virus (MeV) is a highly contagious respiratory virus transmitted via aerosols. To understand how MeV exits the airways of an infected host, we use unpassaged primary cultures of human airway epithelial cells (HAE). MeV typically remains cell-associated in HAE and forms foci of infection, termed infectious centers, by directly spreading cell-to-cell. We previously described the phenomenon in which infectious centers detach *en masse* from HAE and remain viable. Here we investigate the mechanism of this cellular detachment. Via immunostaining, we observed loss of tight junction and cell adhesion proteins within infectious centers. These morphological changes indicate activation of epithelial-to-mesenchymal transition (EMT). EMT can contribute to wound healing in respiratory epithelia by mobilizing nearby cells. Inhibiting TGF-β, and thus EMT, reduced infectious center detachment. Compared to uninfected cells, MeV-infected cells also expressed increased levels of sphingosine kinase 1 (SK1), a regulator of live cell extrusion. Live cell extrusion encourages cells to detach from respiratory epithelia by contracting the actomyosin of neighboring cells. Inhibition or induction of live cell extrusion impacted infectious center detachment rates. Thus, these two related pathways contributed to infectious center detachment in HAE. Detached infectious centers contained high titers of virus that may be protected from the environment, allowing the virus to live on surfaces longer and infect more hosts. This mechanism may contribute to the highly contagious nature of MeV.

**IMPORTANCE:** Measles virus (MeV) is an extremely contagious respiratory pathogen that continues to cause large, disruptive outbreaks each year. Here, we examine a phenomenon that may help explain the contagious nature of MeV: detachment of MeV-infected cells. MeV spreads cell-to-cell in human airway epithelial cells (HAE) to form groups of infected cells, termed “infectious centers”. We reported that infectious centers ultimately detach from HAE as a unit, carrying high titers of virus. Viral particles within cells may be more protected from environmental conditions, such as ultraviolet radiation and desiccation. We identified two host pathways, epithelial-to-mesenchymal transition and live cell extrusion, that contribute to infectious center detachment. Perturbing these pathways altered the kinetics of infectious centers detachment. These pathways influence one another and contribute to epithelial wound healing, suggesting infectious center detachment may be a usurped consequence of the host’s response to infection that benefits MeV by increasing its transmissibility between hosts.

## INTRODUCTION

Measles virus (**MeV**) surpasses other respiratory viruses in transmissibility, but little evidence exists to explain the contagious advantage of MeV. MeV is a morbillivirus in the paramyxovirus family and, like other respiratory viruses, is primarily transmitted via aerosols. In an unvaccinated population, an individual with RSV may spread virus to 3 others, while an individual with MeV may spread virus to 12-18 others (1, 2). Vaccination rates must exceed 92% for herd immunity; however, global vaccination rates have declined to 83% in the past two years, leading to increased incidence of measles outbreaks and elevating MeV’s status to an imminent threat (3, 4, 5).

To model MeV respiratory infections, we use primary human airway epithelial cells (HAE). HAE are derived from human lung donors and are directly seeded onto transwell filters without passage (6). These cultures grow at an air-liquid interface, polarize, and differentiate into a pseudostratified cell layer. HAE recapitulate the airways *in vivo* (7, 8). MeV infects HAE via its epithelial cell receptor, nectin-4 (9, 10). Immune cells, such as dendritic cells or lymphocytes, deliver the virus to nectin-4 on the basolateral side of HAE (11, 12). MeV then replicates in HAE and spreads directly cell-to-cell via intercellular pores (13). MeV typically remains cell-associated in HAE and forms foci of infection, termed infectious centers (shown schematically, **Fig 1A**) (6, 13, 14). Infectious centers are unique to morbilliviruses; viruses outside of this genus do not form infectious centers on HAE (13). Infectious centers are unique from syncytia and multinucleated giant cells as the nuclei remain separate and the plasma membranes between infected cells remain largely intact (6, 13). MeV infectious centers have been observed in HAE, in human tracheal explants (6), and in the nasal passage, trachea, and bronchi of experimentally infected macaques (15, 16).

**Fig 1.**
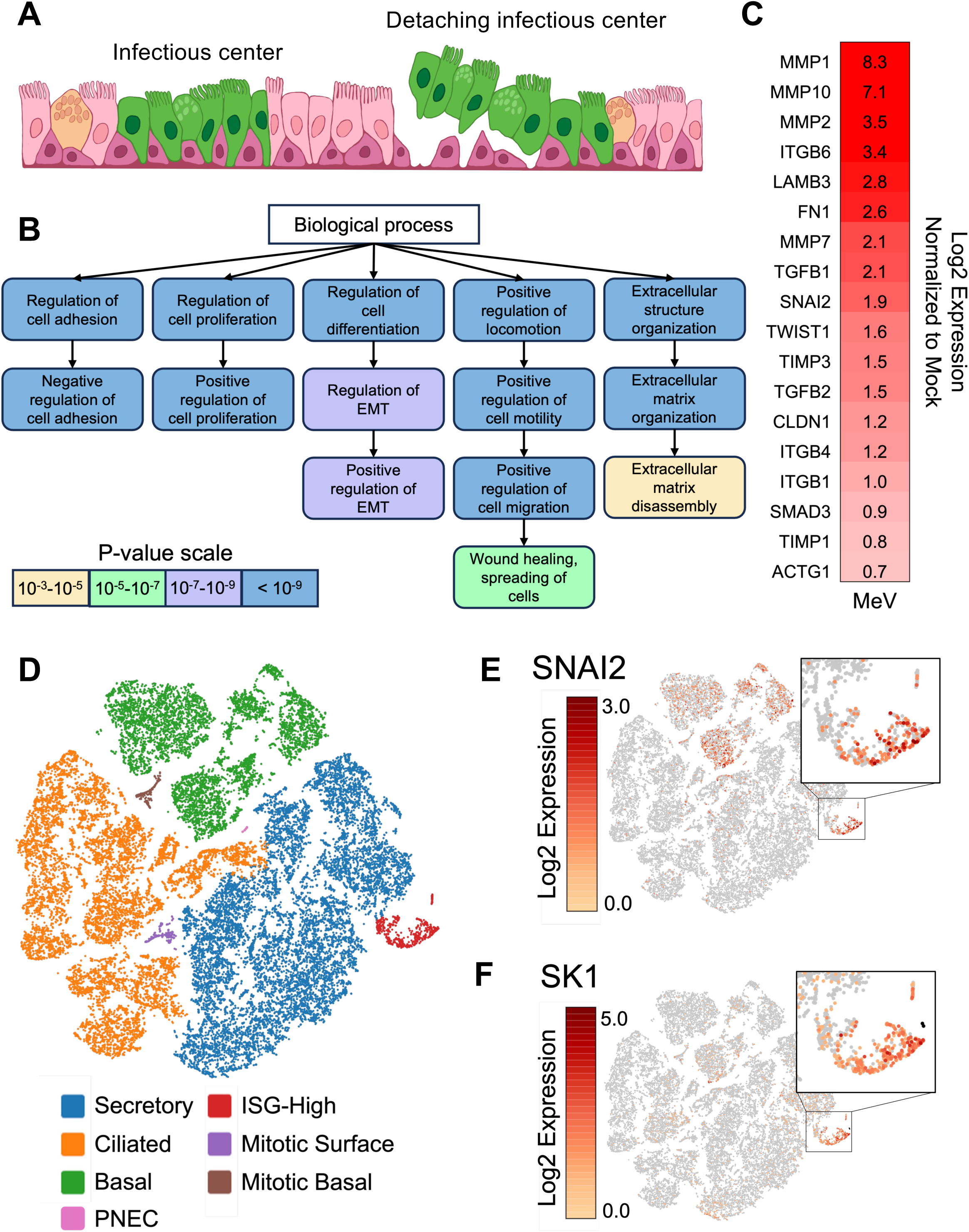
Single-cell RNA sequencing implicates two detachment pathways. (A) Schematic depiction of infectious center formation and detachment. Green cells represent MeV-infected cells. Cell-to-cell spread of MeV over 3-5 days results in an infectious center, left. Beginning around 5 days post-infection, infectious centers begin to detach as a unit from the rest of the epithelial layer, right. (B) Differentially expressed gene lists of MeV-infected HAE were derived from a previously published scRNA-seq dataset (21). Genes with increased mRNA levels in MeV-infected HAE at 3 days post-infection vs. mock-infected HAE were analyzed with biological process gene ontology. Related gene ontology terms are connected with arrows. (C) Heatmap of mRNA expression for EMT-related genes in MeV-infected HAE at day 3 post-infection. Gene expression was normalized to mock-infected HAE and derived from the scRNA-seq dataset. (D) Cells from the scRNA-seq dataset are visualized via t-SNE plot. Cells are clustered by similar gene transcription. Cell types were assigned to clusters using cell type markers described in (21). (E) Cells expressing SNAI2, a transcriptional regulator of EMT, are colored according to mRNA expression. A cluster of cells with high SNAI2 expression is highlighted in the inset. (F) SK1, which promotes LCE, is also expressed in this cell cluster (see inset).

Most respiratory viruses leave airway epithelial cells via apical budding, exocytosis, or cell lysis. Many enveloped RNA respiratory viruses (e.g., IAV (17), RSV (18), and SARS-CoV-2 (19)) utilize apical budding at the host cell’s plasma membrane to exit airway epithelial cells. Non-enveloped viruses, like rhinovirus, exit via vesicles (exocytosis) or by bursting the cell open (cell lysis) (20). MeV, an enveloped virus, was assumed to exit HAE via apical budding; however, we found that MeV infection uniquely leads to dislodging of infectious centers containing concentrated virus (**Fig 1A**) (21). Evidence for MeV-induced cell detachment is also found in monkey epithelial cultures, *in vivo* monkey experiments, and human observation studies (15, 22, 23). The cellular signals that promote IC detachment are not known; however, cells of detached infectious centers are metabolically active and do not display markers of cell death pathways, suggesting live cell extrusion pathways are involved (21, 23).

Here, we investigate the mechanism of infectious center detachment. As will be shown, single-cell RNA sequencing (21) suggests epithelial-to-mesenchymal transition (EMT) is activated within infectious centers. Airway epithelial cells are typically quiescent and nonmigratory. However, certain circumstances necessitate epithelial cells to divide and migrate. EMT is initiated by diverse stimuli to dissolve cell-cell connections and promote mobilization. EMT is involved in wound repair, tumor metastasis, and response to viral infection (24, 25, 26). Importantly, EMT encompasses a spectrum of intermediate cell types.

EMT signaling associates with live cell extrusion (LCE), a pathway driven by sphingosine kinase 1 (SK1). SK1 phosphorylates sphingolipids, which are cell signaling molecules that localize to membranes. In infected cells, increased SK1 activity drives pro-survival signaling that results in more virus production (27, 28, 29, 30, 31, 32). In addition, downstream signaling of SK1 directs neighboring cells to contract their actomyosin, creating a luminal force on the cell to be extruded (33, 34). Cell extrusion is important for maintaining homeostasis within epithelia and can be activated by several stimuli, including EMT, apoptosis, overcrowding, or loss of cell polarity (35, 36, 37). LCE and EMT signaling pathways overlap and each can activate the other. Here, we show that both pathways are active within infectious centers and contribute to detachment.

## RESULTS

### Single cell RNA-sequencing implicates two detachment pathways

To investigate what pathways may be responsible for infectious center detachment, we probed our previously published single-cell RNA sequencing (scRNA-seq) dataset (21). In this dataset, HAE from 10 human donors were mock-infected or infected with a MeV expressing a GFP reporter protein (MeV-GFP). At multiple timepoints post-infection, samples were sorted via fluorescence-activated cell sorting (FACS) to enrich the GFP+ population before performing scRNA-seq. Lists of genes differentially expressed between mock and MeV infected (both GFP+ and GFP-) HAE were analyzed by gene ontology (**Fig 1B**). Many gene ontology terms from this analysis pointed to EMT processes, including “positive regulation of EMT”. Enrichment of gene products such as TGFB1, SNAI2, and metalloproteinases (MMPs) are emblematic of EMT (**Fig 1C**).

EMT-related mRNAs were notably increased in a distinct cluster of cells, termed “ISG-high” (**Fig 1D and 1E**). This cluster was exclusively observed in MeV-infected HAE cultures (**S1A Fig**), contained ∼80% GFP+ and ∼20% GFP-cells, and the cells expressed unique transcriptomes that did not align with known HAE cell types (**Fig 1D**). Previously, we erroneously dubbed these cells as “interferon-high cells” due to elevated interferon-stimulated gene (ISG) transcripts (21). However, further analysis revealed that expression of these ISGs was not interferon-mediated (38). As such, we renamed these cells “ISG-high cells” and continue to investigate their characteristics. We could not identify what cell type the ISG-high cells were before they were infected possibly because of the transcriptional impact of EMT.

Notably, ISG-high cells had elevated SNAI2 transcripts, a transcriptional regulator of EMT (**Fig 1E and S1B Fig**). Further investigation of the transcriptional profiles of ISG-high cells revealed a potential second detachment pathway: live cell extrusion (LCE). LCE is induced by sphingosine kinase 1 (SK1), one of the top enriched gene products within the ISG-high cluster (**Fig 1F and S1C Fig**). In this manuscript, we will separately consider EMT and LCE for infectious center detachment.

### EMT is activated within infectious centers

We evaluated the role of EMT in MeV infection. Quantitative Real-Time PCR (qRT-PCR) confirmed EMT-related gene transcripts are elevated in MeV-infected cultures (**S2A Fig**). In addition, TGF-β, a potent inducer of EMT, was secreted in higher concentrations following MeV-infection (**S2B Fig**). EMT is characterized by a loss of epithelial cell markers (39, 40, 41, 42). Indeed, immunostaining of infectious centers at 3 days post-infection and 7 days post-infection with MeV-mCherry showed a loss of β-catenin (**Fig 2A**), ZO-1 (**Fig 2B**), and E-Cadherin (**Fig 2C**). In all cases, the loss of the markers was more pronounced toward the center of the infectious center. Degradation increased over time and ∼60% of the infectious center area had loss of the indicated junctional protein by day 7 (**Fig 2D**). Additionally, cells of infectious centers may have altered apical-basal polarity, as evidenced by an atypical pattern of expression of the apically targeted ezrin protein (arrow; **S2C Fig**). These results demonstrate EMT activation in MeV-infected HAE.

**Fig 2.**
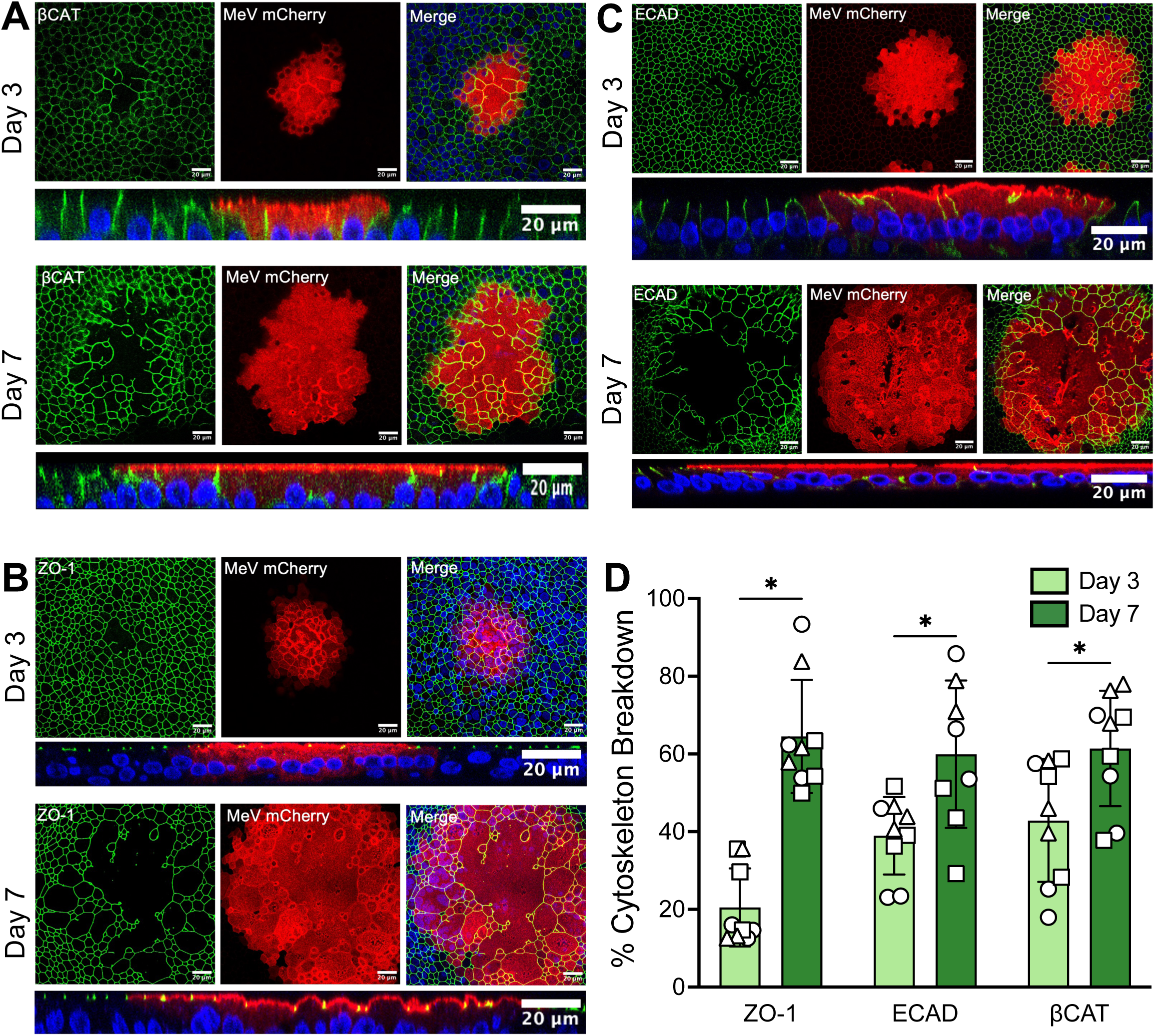
EMT is activated within infectious centers. HAE infected with MeV-mCherry were fixed at 3- and 7-days post-infection. Cultures were immunostained for βCAT (A), ZO-1 (B), or ECAD (C). Confocal images of 3 infectious centers were collected per donor (n=3). Representative images are shown. Corresponding z-stacks are shown below each image. Blue = DAPI. (D) Percent cytoskeleton breakdown was calculated by dividing total area of breakdown for each cytoskeletal protein by the total area of the infectious center. Multiple t-tests were performed. * p<0.05.

### EMT contributes to infectious center detachment

Infectious center detachment can be tracked using live cell fluorescence microscopy. Images were collected daily post-infection and the timepoint at which each infectious center detached was documented (**Fig 3A**). These data were used to calculate a probability of detachment. To determine the impact of EMT on infectious center detachment, we treated MeV-infected HAE with a TGF-β inhibitor, SB431542, which blocks downstream transcriptional activation of TGF-β (43, 44). With TGF-β inhibited, infectious centers detached at a slower rate than vehicle-treated controls (**Fig 3B**). Titers (**Fig 3C**), infectious center number (**Fig 3D**), infectious center area (**Fig 3E**), and cytotoxicity (**Fig 3F**) were not statistically different between the treatment and control groups, suggesting that TGF-β inhibition slows IC detachment without impacting viral replication or cell-to-cell spread.

**Fig 3.**
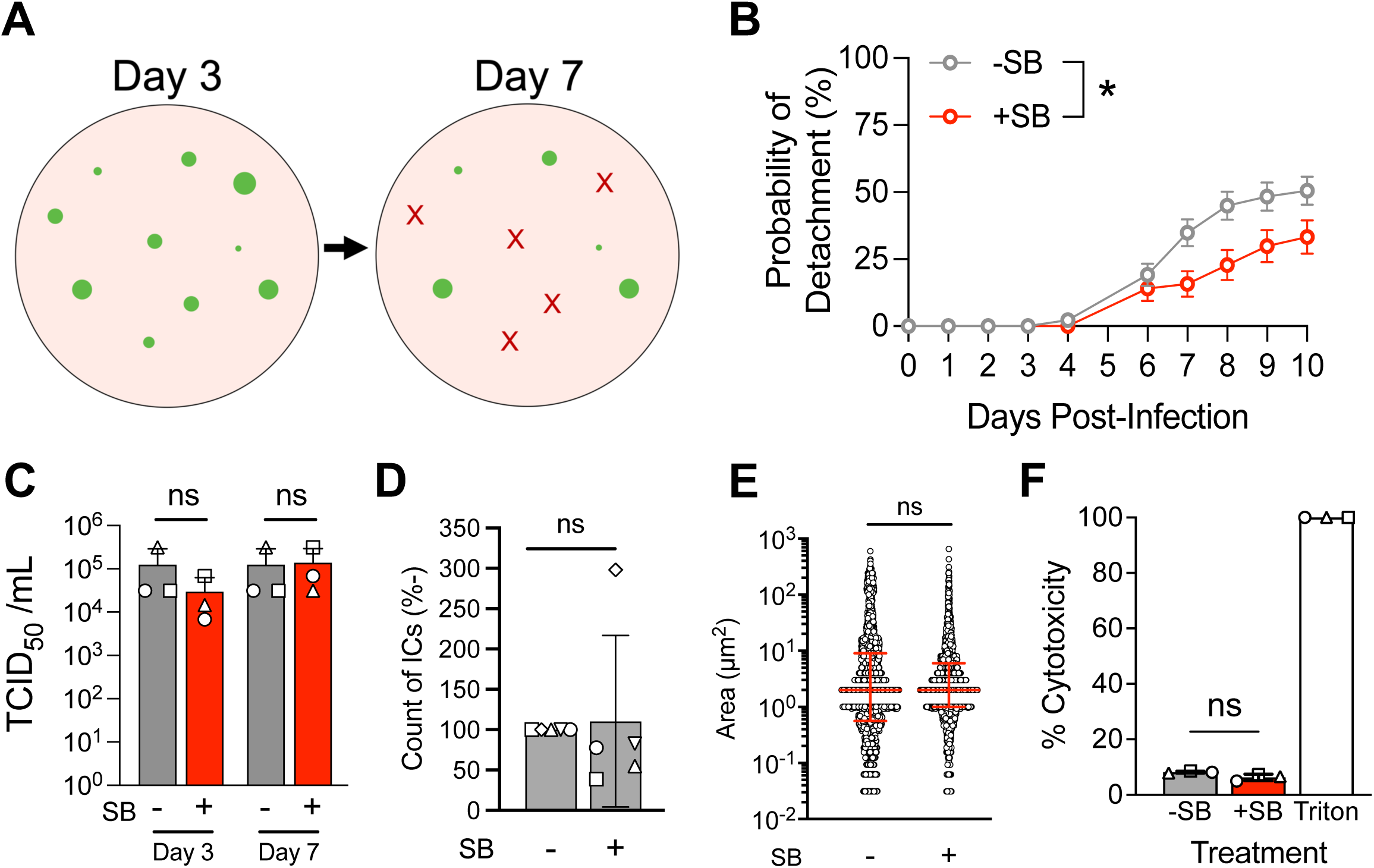
EMT contributes to infectious center detachment. (A) Graphic representation of infectious center detachment assay. Images are taken each day post-infection. Individual infectious centers (green dot) are tracked until they detach (represented by a red X). The day post-infection at which each infectious center detaches is recorded. (B) HAE were treated with either SB431542 (SB) or a vehicle control (DMSO) for 24 hours before being infected at an MOI of 1 (MeV-GFP). Data from the detachment assay are graphed as a probability of detachment (i.e., percent of infectious centers detached at each time point). A Gehan-Breslow-Wilcoxon test was performed and indicated that there was a statistical significance between the two curves. * p<0.05. (C) Whole culture lysates, which may include detached infectious centers in the apical space, were collected from MeV-infected HAE (with either SB treatment or vehicle) at the indicated timepoint for titration via TCID_50_. A one-way ANOVA with multiple comparisons was performed. ns = not significant. (D) Live fluorescent images were collected at 3 days post-infection. Numbers and size (E) of infectious centers were measured with ImageJ. Counts were normalized to vehicle controls and displayed as a percentage. Shapes indicate individual donors (n=5). Student’s t-test. (E) Dots indicate individual infectious centers. Median ± interquartile range. Student’s t-test. (F) Cytotoxicity of SB treatment was determined via LDH assay at 3 days post-infection with MeV. SB and DMSO treated cells had comparable cytotoxicity when normalized to Triton X-100 treated cells (+ control). One-way ANOVA.

### LCE is activated within infectious centers

EMT can lead to LCE. Once a cell is fated for extrusion, sphingosine, a cellular lipid, is phosphorylated by sphingosine kinase 1 (SK1) to produce sphingosine-1-phosphate (S1P) (**Fig 4A**). S1P binds its receptor, S1PR2, on neighboring cells, signaling for the activation of the Rho pathway. As a result, a ring of cells surrounding the fated cell will contract their actomyosin, creating an apical force to expel the cell (33, 34, 35, 37). Transcripts for *SK1*, Rho pathway genes (*RHOA*, *RHOB*, and *RHOD*), and genes responsible for actomyosin contraction (*MYH9* and *ROCK1*) (37, 45) were elevated within the ISG-high cell cluster of our scRNA-seq dataset (**Fig 4B**). Immunostaining revealed altered SK1 localization within the cells of infectious centers (**Fig 4C and 4D**). Specifically, SK1 was most abundant at the nucleus (arrows; **Fig 4D**). Together, these data suggest SK1 is expressed in MeV-infected cells and initiates LCE signaling.

**Fig 4.**
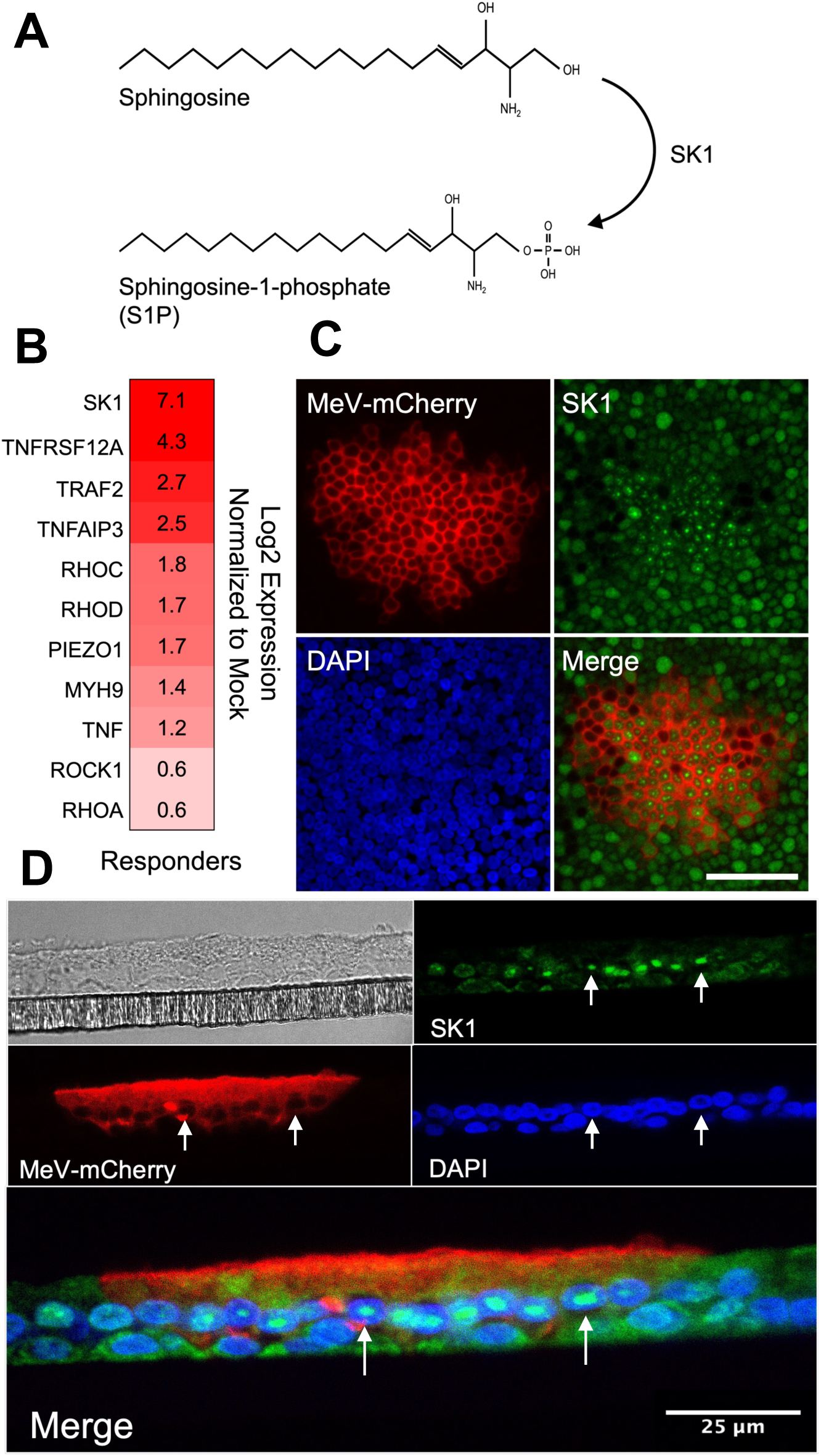
LCE is activated within infectious centers. (A) Diagram of SK1’s function. SK1 phosphorylates sphingosine to make S1P. (B) Heatmap of LCE-related gene expression in ISG-high cells at 3 days post-infection. Shade of red indicates increased fold change over mock-infected cells. (C) *En face* immunostaining of SK1 in HAE infected with MeV-mCherry at 3 days post-infection. Scale bar = 50 μm. (D) Immunostained sections of MeV-mCherry infected HAE at 3 days post-infection. Arrows indicate cells with SK1 staining localized to the nucleus. Scale bar = 25 μm.

### LCE contributes to infectious center detachment

Using the infectious center detachment assay described in Figure 3, we evaluated the impact of an SK1 inhibitor on infectious center detachment. Treatment with the SK1 inhibitor, SKI-II, delayed infectious center detachment (**Fig 5A**). SKI-II treatment did not cause a statistically significant change in titers (**Fig 5B**), infectious center number (**Fig S3A**), or infectious center area (**Fig S3B**) of MeV-infected HAE. To exacerbate LCE, MeV-infected HAE were treated with exogenous S1P. The probability of infectious center detachment increased and infectious centers detached faster (**Fig 5C**), despite viral titers (**Fig 5D**), infectious center number (**Fig S3C**), infectious center area (**Fig S3D**), and cytotoxicity (**Fig S3E**) remaining statistically indistinguishable to vehicle-treated controls. These results demonstrate infectious center detachment utilizes LCE pathways.

**Fig 5.**
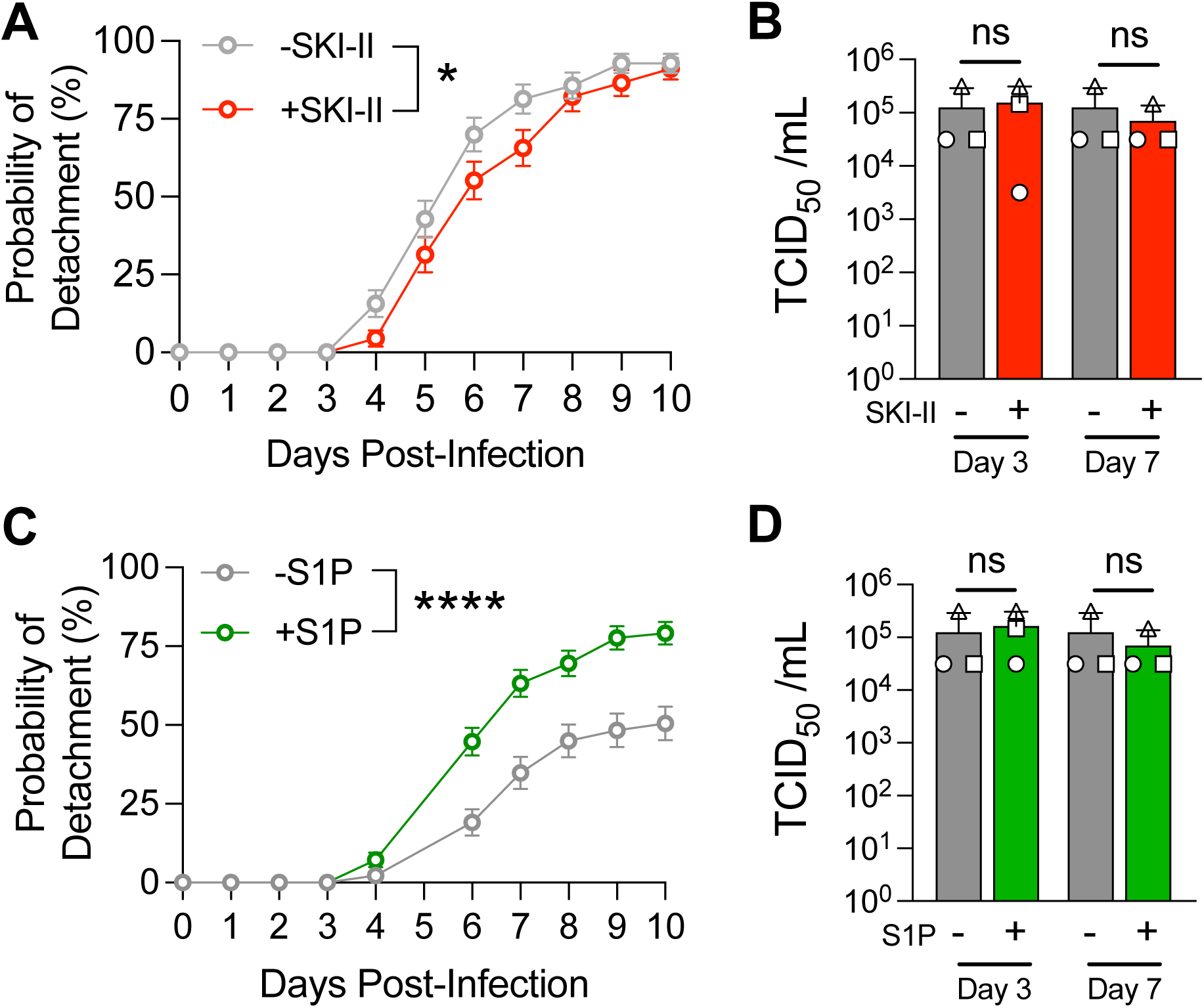
LCE contributes to infectious center detachment. (A) An SK1 inhibitor (SKI-II) or (C) exogenous S1P was applied 1-day post-infection with MeV. The infectious center detachment assay was performed as described in Fig 3. Gehan-Breslow-Wilcoxon tests were performed and indicated that there were statistical significances between the curves. * p<0.05; **** p<0.0001. (B) Cell lysates were collected from MeV-infected HAE treated with SKI-II or (D) S1P at the indicated timepoint for titration via TCID_50_. A one-way ANOVA with multiple comparisons was performed. ns = not significant.

## DISCUSSION

Here, we showed that two overlapping, wound healing pathways are active within the cells of infectious centers. Transcriptional regulators of EMT and LCE are elevated in MeV-infected cells. Physical evidence of EMT and LCE was observed via immunostaining and microscopy. Additionally, because infectious center detachment could be delayed by inhibiting EMT and LCE and increased using an LCE agonist, we concluded that both pathways contribute to infectious center detachment. Although not shown, we induced EMT in HAE with exogenous TGF-β followed by MeV infection to determine if infectious center detachment increased. However, the results were uninterpretable because infection of epithelial cells primed for EMT results in widespread syncytial-like, multi-nucleated giant cells instead of discreet infectious centers. As we previously reported, infectious center formation requires proper epithelial differentiation (6). As primary epithelial cell culture has become more accessible, cell detachment has been observed in several virus infections of epithelia. This includes astrovirus (46), rotavirus (47), and enterovirus A71 (48). Both EMT and LCE are candidate mechanisms for cell detachment in these infections and are potentially a common result of epithelial cell viral infection.

EMT and LCE pathways have important roles in maintaining homeostasis of the epithelial layer and their functions often overlap (35, 45, 49, 50). LCE activates in cases of cell overcrowding (34, 36, 51), removing damaged cells following injury (51), or following apoptotic signaling (37, 52, 53, 54). EMT is a key tool for wound healing and is active in chronic lung diseases like idiopathic pulmonary fibrosis (55). In EMT, cells can gain actomyosin contractility, a key mechanism for LCE (41). Transcriptional regulators of EMT, like SNAI1, impact RhoA signaling and actomyosin contractility (45). Wee et al. (45) showed that cell-cell adhesion loss works cooperatively with actomyosin contraction to cause cell extrusion. In a complementary manner, SK1 promotes EMT by degrading cell-cell adhesions, like E-cadherin, and activating the focal adhesion kinase pathway, which includes MMPs (56, 57). Inhibiting SK1 blocks signaling of two EMT activators, NF-κB and TGF-β, whereas exogenous S1P treatment induces EMT (58, 59). Notably, SK1 and SNAI2 coexpress within human tumors (60). In fact, LCE and EMT appear to be so intertwined that one does not exist without the other. We postulate that, in the context of MeV infection of HAE, EMT and LCE activate simultaneously and cooperate to expel infectious centers. EMT likely promotes infectious center detachment by altering cell-cell adhesions, while LCE mechanically promotes detachment via actomyosin contraction.

SK1 is involved in several viral infections because of its role in activating cell pro-survival signaling (29, 30, 61, 62, 63, 64). A stressed cell will accumulate sphingosine in its plasma membrane, marking the cell for apoptosis; however, in the presence of increased SK1 expression, sphingosine is phosphorylated by SK1 to create S1P, a signal for cell survival (37, 65, 66, 67). S1P prolongs the life of an infected cell and increases viral replication as the virus has more time to propagate. Viral replication is associated with SK1 expression during infection with respiratory syncytial virus (32), human cytomegalovirus (68), and influenza virus (28). In cases of viral infections, SK1 expression is induced via TNF-α signaling and regulated by the ERK/AKT pathway (69, 70, 71). MeV may also take advantage of SK1’s cell survival phenotype to increase virion production. Indeed, several genes of the TNF-α pathway were upregulated in our scRNA-seq ISG-high cells (**Fig 4B**).

We observed SK1 expression in the nucleus of infected cells (**Fig 4C and 4D**). While SK1 is more commonly found in the cytoplasm, nuclear localization is seen in human lungs with idiopathic pulmonary fibrosis (72) and SK1 translocation to the nucleus is induced with estrogen treatment of breast cancer cells (73). While the role of nuclear SK1 is still unclear, sphingosine kinase 2 is known to alter gene expression in the nucleus and may render cells more resistant to apoptosis (74). In addition, MeV replication was previously shown to be sensitive to SK1 inhibition in HEK 293 cells (30). In that study, NF-κB expression was necessary for MeV replication and inhibiting SK1 equally inhibited NF-κB. In contrast, we did not find that SKI-II treatment affected viral titers in HAE. This may be due to differences in cell types used since HAE are much more resilient than HEK 293 cells to viral infections and are more likely to survive. Importantly, immortalized tumor cells are likely to be inherently skewed toward a mesenchymal phenotype even if they were originally derived from an epithelial cell. Thus, unpassaged, primary epithelial cells may be better suited for studying viral induced EMT and LCE.

EMT might also explain why infectious centers stop growing. Our recent work determined that cell-cell spread of MeV is not restricted by interferon-induced innate immune signaling (38). EMT causes the disassociation of cell-cell junctions, including adherens junctions (24). The MeV receptor nectin-4 is localized in adherens junctions and is required for cell-cell spread (13). The loss of adhesion caused by EMT could contribute to halted infectious center growth. This sequestration of infectious centers from the rest of the epithelia may also explain why the cells of infectious centers maintain their connections to one another while simultaneously losing their connection to neighboring cells. While we do not have an answer for why cells of infectious centers remain attached to one another after they are dislodged, we have two hypotheses: 1) Cells with similar actin densities group together (75): as MeV infection degrades actin within infectious centers, the surface tension of the cellular membranes of infectious center cells may decrease, distinguishing these cells from the uninfected cells around them. 2) While mesenchymal-like cells do not integrate well within tissues during migration, they can form transient cell-cell adhesions with each other and nearby stationary cells (76). One such connection could be formed during the transfer of MeV factors through intercellular pores that form during cell-cell spread. Future studies identifying cell-cell adhesions that persist in MeV-infected cells will be key in determining the bottlenecks that impact infectious center growth and detachment.

We identified a group of “ISG-high cells” in our scRNA-seq that had high transcripts of ISGs as well as EMT and LCE markers. We postulate that these cells are either within infectious centers or are uninfected cells near the periphery of infectious centers (**Fig S1A**). These cells are responding to the infection, seemingly by increasing antiviral gene transcripts. In a recent publication, we studied interferon stimulation in HAE in the context of MeV infection and determined interferons are not produced at detectable levels (38). While the mystery of what initiates ISG transcription remains, we believe the name “ISG-high cells” more accurately describes this cluster than the previously reported “interferon high cells”(21). Importantly, we are not suggesting that ISG-high cells are a new epithelial cell type, but cells with an altered transcriptional profile following MeV infection. ISG-high cells were likely secretory or ciliated cells before MeV infection, but we cannot be sure because those cell type markers are not detectable in our samples. Understanding the influences that push cells towards the ISG-high cell transcriptome profile is of interest.

In summary, our results describe two, related pathways by which MeV extrudes groups of virus-laden cells. Infectious center detachment was altered by inhibitors and agonists of EMT and LCE, which supports our conclusion that MeV exits HAE via this strategy. Damage to the cell during MeV infection likely initiates these pathways and infectious center detachment is an unintended consequence that gives MeV an advantage. While vaccination remains the best preventative measure for MeV, understanding MeV infectious center detachment is vital for investigating the contagious nature of MeV and may prove helpful in healthcare settings.

## MATERIALS AND METHODS

### Ethical statement

The University of Iowa *In Vitro* Models and Cell Culture Core prepared the well-differentiated primary cultures of human airway epithelia (HAE) used in these studies. These cultures are comprised of cells from autopsy, discarded tissue, or surgical specimens. All human subject studies were conducted with approval from the University of Iowa Institutional Review Board. We do not receive identifiable information.

### Human airway epithelial cells

HAE were cultured and maintained as previously described (6, 7). Briefly, cells from trachea and bronchus epithelia are seeded onto collagen-coated, polycarbonate transwell inserts (0.4 μm pore size, surface area = 0.33 cm^2^; Corning Costar; #3413) following enzymatic disassociation. Cells were submerged in Ultroser G (USG) medium at 37°C and 5% CO_2_. After 48 hours, apical USG was removed and cells were incubated at the air-liquid interface and allowed to polarize and differentiate. All HAE cultures used for the experiments in this study were >3 weeks old and maintained a transepithelial electrical resistance of at least 500 ν⋅μm^2^.

### Bioinformatic analyses

The scRNA-seq dataset was previously described (21) and can be accessed in the GEO with accession number GSE168775. In brief, MeV-infected or mock-infected HAE (pooled from 10 human donors) were dissociated using TrypLE and sorted by fluorescence-activated cell sorting to separate GFP- and GFP+ cells for scRNA-seq. ∼30,000 cells make up the dataset. Gene ontology analysis was performed using GOrilla (77, 78). Violin plots, t-SNE plots, and heatmap data were generated in Loupe Browser version 6.4.1. Heatmaps were visualized with the R package gplots version 3.1.3.

### Measles virus production

Recombinant MeV-GFP and MeV-mCherry viruses are derived from the IC323 strain and were produced as previously described (79). Briefly, Vero cells that stably express human SLAMF1 (80) were maintained in Dulbecco modified Eagle medium (DMEM; Thermo Fisher Scientific) supplemented with 5% newborn calf serum (NCS; Thermo Fisher Scientific) and penicillin-streptomycin (100 mg/mL; Thermo Fisher Scientific). Vero-hSLAM cells were infected with either MeV-GFP for MeV-mCherry at an MOI of 0.01 for 2 hours. 2 days post-infection, cells were collected and subjected to several freeze/thaw cycles to induce cell membrane rupture and virus release. Virus was titrated using TCID_50_ assays with Vero-hSLAMF1 cells. We have not detected differences in MeV-GFP and MeV-mCherry infections. We selected MeV-GFP or MeV-mCherry for our experiments due to their compatibility with reagents and instrumentation.

### Infection of HAE

HAE cultures in this study were infected with MeV as previously published (13, 14). Briefly, HAE cultures were inverted to expose the basolateral surface. 50 μL virus inoculum, containing MeV virus and serum-free opti-MEM (Thermo Fisher Scientific; #31985062) was placed on the basolateral side of the cells. Virus inoculum was incubated on HAE for ∼4 hours at 37°C and 5% CO_2_. After the infection period, the inoculum was removed and the cultures were placed back into an upright position with medium in the basolateral chamber.

### Immunostaining and confocal microscopy

All cells prepped for immunofluorescence were fixed with 2% paraformaldehyde for 15 minutes followed by washing with 1X PBS and permeabilization with 0.2% Triton X-100 in Superblock (Thermo Fisher Scientific; #37580) for 1 hour. All primary antibodies were used at 1:100 dilution in Superblock overnight at 4°C. After overnight incubation, cells were washed 3x with 1X PBS and incubated with fluorescently labelled secondary antibodies at a 1:1000 dilution in Superblock buffer for 1 hour at RT, gentle rocking. HAE filters were cut from the transwell using a razor blade and mounted onto glass microscope slides, covered with VECTASHIELD Mounting Medium with DAPI (Vector Laboratories; #H-1200-10) and a glass coverslip overtop. Coverslips were sealed with clear nail polish. For confocal analysis, a Leica SP8 STED confocal microscope (Leica Microsystems) was used. Image processing was performed using ImageJ2 version 2.3.0. Primary antibodies used: Beta-catenin (Thermo Fisher Scientific; #13-8400), ZO-1 (Thermo Fisher Scientific; #33-9100), E-cadherin (Cell Signaling Technology; #3195), Ezrin (Cell Signaling Technology; #3145), SK1 (Invitrogen; #PA5-28584). Secondary antibodies used: Goat Anti-Rabbit IgG Highly Cross-Adsorbed (H+L) Alexa Fluor 568 (Invitrogen; #A-11036), Goat Anti-Mouse IgG Highly Cross-Adsorbed (H+L) Alexa Fluor 546 (Invitrogen; #A-11030).

### Quantification of cytoskeleton breakdown

Confocal images of 3, randomly selected infectious centers were collected from 3 human donors for BCAT, ZO-1, and ECAD. Infectious center area was measured using the freehand tool in ImageJ2 version 2.3.0. Adhesion disruption area was determined by accumulating the area measurements of all protein degradation within the infectious center. The data are plotted as a percentage: (total area of degradation) / (total area of infectious center) x 100.

### Quantitative reverse transcriptase polymerase chain reaction (qRT-PCR)

At 3 dpi, mock- and MeV-infected cell lysates were collected with TRIzol reagent (Thermo Fisher Scientific; #15596026). Cellular RNA was isolated using the Direct-zol RNA Miniprep kit (Zymo Research; #R2052). After isolation, RNA was transcribed into cDNA with the High-Capacity RNA-to-cDNA kit and the manufacturer’s instructions (Thermo Fisher Scientific; #R2052). qPCR of *MMP1*, *MMP10*, *TGFB1*, *SNAI2*, and *SMAD3* was accomplished using primers, *Power*SYBR Green Real-Time PCR Master Mix (Thermo Fisher Scientific; #4367659), and a QuantStudio 7 Pro (Applied Biosystems). The housekeeping gene used was *SFRS9*. Primer sequences can be found in Table S1.

### TGF-β ELISA

Apical washes (100 µl Opti-MEM) and basolateral media (100 µl USG medium) were collected from mock- and MeV-infected HAE. Secretions were measured in duplicate for TGF-β protein content using a Human TGF-beta 1 ELISA Kit (RayBiotech; #ELH-TGFb1-1) and following the manufacturer’s protocol. ELISAs were read on XXX.

### Drug treatments

SKI-II (MedChemExpress; #312636-16-1) and SB431542 (Tocris; #1614) were diluted to 20 μM with USG medium. Recombinant S1P (Tocris; #1370) was diluted to 10 μM with USG medium. Treatments were placed in the basolateral chamber of the HAE cultures. Equivalent volume of DMSO (4 μl) was diluted in USG medium for vehicle control cultures. SKI-II and S1P treatments began 1-day post-infection. SB431542 treatment began 1 day pre-infection. Treatments remained on the cells for the length of the experiment and media/treatments were refreshed every 2 days.

### Live-image microscopy and infectious center analyses

Image acquisition was performed using an All-in-one-Fluorescence Microscope BZ-X800E (Keyence Corporation of America) with a 4X objective. Images from each culture were arranged sequentially by time post-infection. A blinded individual tracked individual infectious centers over the time course of the experiment and determined which day post-infection the infectious center detached. These data were converted to binary and were graphed as a probability of detachment.

### Titration of MeV

MeV-infected HAE lysates for titration analysis were scraped and collected in serum-free medium and 10-fold serially diluted. Serial dilutions were plated in the 96-well flat-bottom plate containing Vero-hSLAM cells. 3 days post-infection, cells were titrated using TCID_50_/mL analysis.

### LDH cytotoxicity colormetric assay

HAE cultures were infected and supernatants from the basolateral chamber were collected. Each cultures received fresh medium after collection. LDH release was detected following the manufacturer’s protocol (BioVision; #K311). Briefly, supernatant and reaction mixture were mixed 1:1 and incubated in a 96-well Maxi-Sorp Immunoplate (Thermo Fisher Scientific; #439454). After 30 minutes at room temperature, absorbance was measured at 490_OD_. Percent cytotoxicity was measured using the following formula: (test sample – low control) ÷ (high control – low control). Mock-infected supernatants served as the low controls. High control samples were obtained by incubating cells with media containing 0.1% Triton X-100 for 30 minutes at 37°C, followed by media collection. This method allows for complete cell death, and therefore maximum LDH release, to occur.

### Sectioning HAE

HAE were fixed with 2% paraformaldehyde for 15 minutes at room temperature. Filters were excised from the transwell, submerged in Scigen Tissue-Plus O.C.T. Compound (Thermo Fisher Scientific; #23-730-571), and frozen. 10 μm sections were collected with a Microm Cryostat HM505E (Leica) and immunostained following the protocols listed above.

### Statistics

Statistical tests listed in the figure legends were performed in GraphPad Prism version 10.1.1. Alpha values were set at 0.05. Unless otherwise indicated, numerical data are presented as mean ± standard deviation (SD).

## ACKNOWLEDGEMENTS

We thank Roberto Cattaneo for providing fluorescent MeVs and guidance. Additionally, we thank Jennifer Bartlett for her thoughtful review of this manuscript. We acknowledge the support of the University of Iowa Central Microscopy Research Facility, the Genomics Division of the Iowa Institute of Human Genetics, and the In Vitro Models and Cell Culture Core.

## AUTHOR CONTRIBUTIONS

Conceptualization: Camilla E. Hippee, Brajesh K. Singh, Patrick L. Sinn.

Data curation: Camilla E. Hippee.

Formal Analysis: Camilla E. Hippee, Lorellin A. Durnell, Justin W. Kaufman.

Funding Acquisition: Camilla E. Hippee, Patrick L. Sinn.

Investigation: Camilla E. Hippee, Lorellin A. Durnell, Justin W. Kaufman, Eileen Murray.

Visualization: Camilla E. Hippee, Lorellin A. Durnell, Justin W. Kaufman.

Writing – original draft: Camilla E. Hippee.

Writing – review & editing: Camilla E. Hippee, Lorellin A. Durnell, Patrick L. Sinn.

## FIGURE LEGENDS

**Fig S1. Characterization of ISG-high cells.** (A) The t-SNE plot from Fig 1 with GFP status overlayed. GFP+ indicates infected cells, GFP-indicates uninfected cells from MeV-infected HAE. Mock indicates cells mock infected. The box highlights the ISG-high cell cluster. The adjoining graph identifies the division of ISG-high cells that are GFP+ or GFP-. (B) Violin plot of SNAI2 expression across cell clusters. (C) Violin plot of SK1 expression across cell clusters.

**Fig S2. qRT-PCR confirmation of scRNA-seq.** (A) scRNA-seq results were confirmed with qRT-PCR. Transcript changes of MeV-infected HAE are shown as a fold change over mock-infected cells. One-way ANOVA was performed. * p<0.05, ** p<0.01. (B) Apical (open shapes) and basolateral (filled shapes) secretions were collected from mock- and MeV-infected HAE at 3 days post-infection. Protein content was measured using a TGF-β ELISA kit. Shapes indicate distinct donors (n=3). Paired Student’s t-test; * p<0.05. (C) HAE infected with MeV-mCherry were fixed at 7 days post-infection and immunostained for ezrin. A z-stack image is shown below. The arrow points to ezrin staining that has been disturbed within the infectious center. Blue = DAPI. Scale bar = 50 μm.

**Fig S3. Impact of LCE treatments on infection kinetics.** Live fluorescent images of (A-B) SKI-II treated or (C-D) S1P treated HAE were collected at 3 days post-infection with MeV. Numbers (A and C) and size (B and D) of infectious centers were measured with ImageJ. Counts were normalized to vehicle controls and displayed as a percentage. Shapes indicate individual donors (n=4 in A; n=5 in C). Student’s t-test; ns = not significant. (B and D) Dots indicate individual infectious centers. Median ± interquartile range. Student’s t-test. (E) Cytotoxicity of SB treatment was determined via LDH assay at 3 days post-infection with MeV. SKI-II, S1P, and DMSO treated cells had comparable cytotoxicity when normalized to Triton X-100 treated cells (+ control). One-way ANOVA.

**S1 Table.**
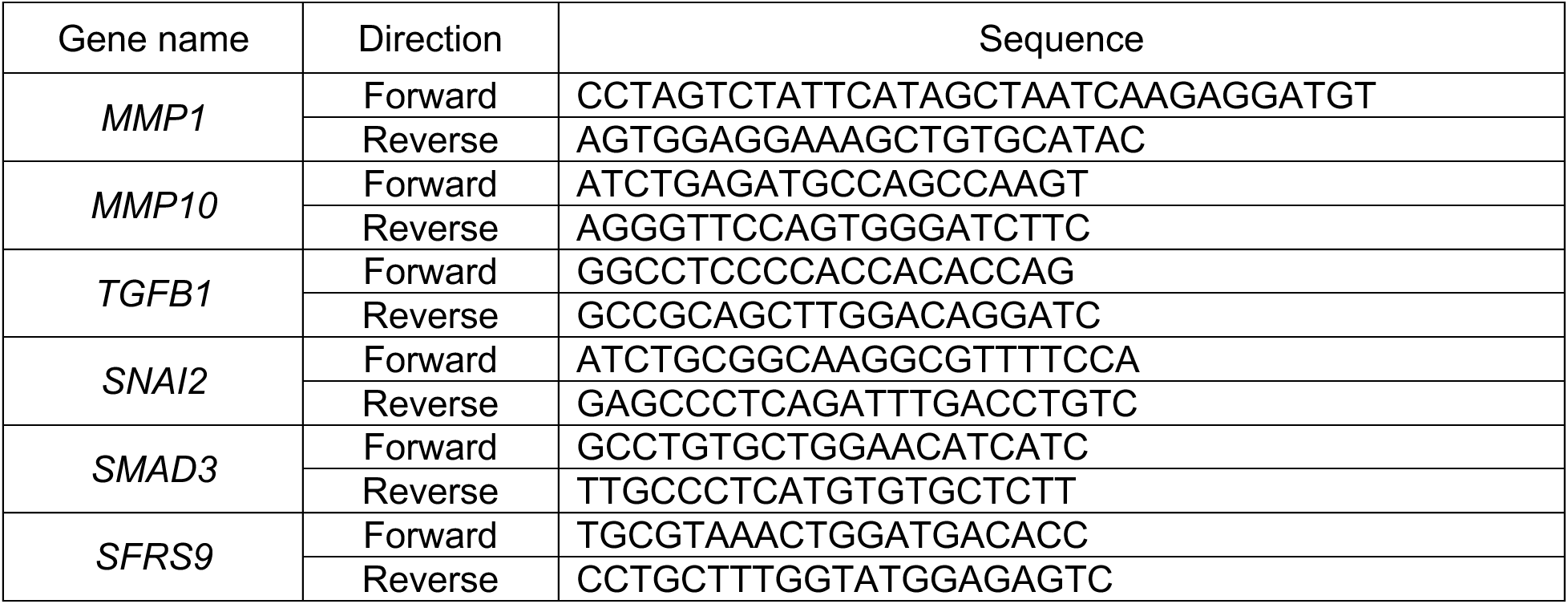
Primer sequences for qRT-PCR analysis.

