## Supplemental Figures for "Measles virus co-opts epithelial-to-mesenchymal transition and live cell extrusion to exit human airway epithelia"

Fig S1. Characterization of ISG-high cells.

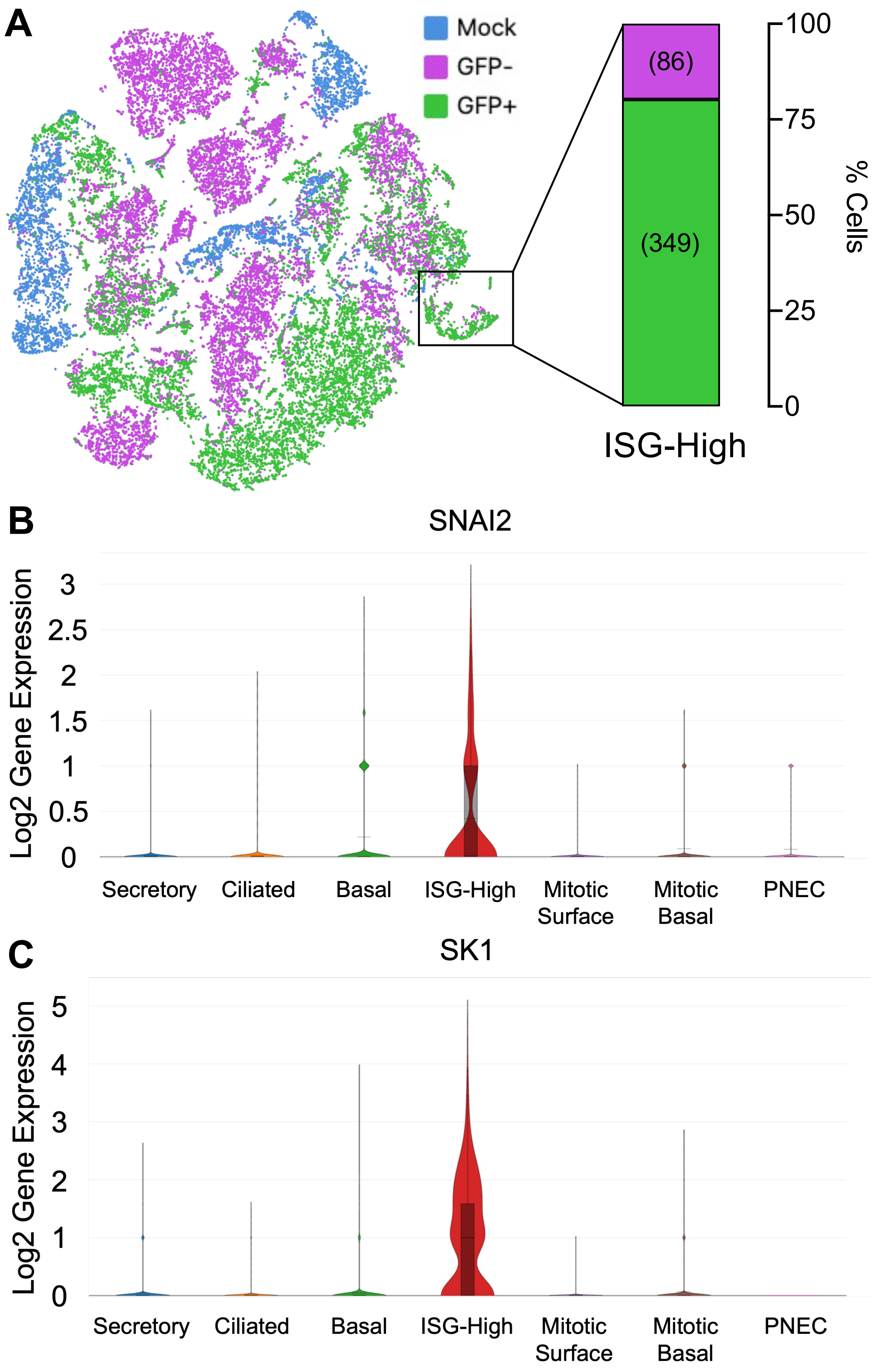

Fig S2. qRT-PCR confirmation of scRNA-seq.

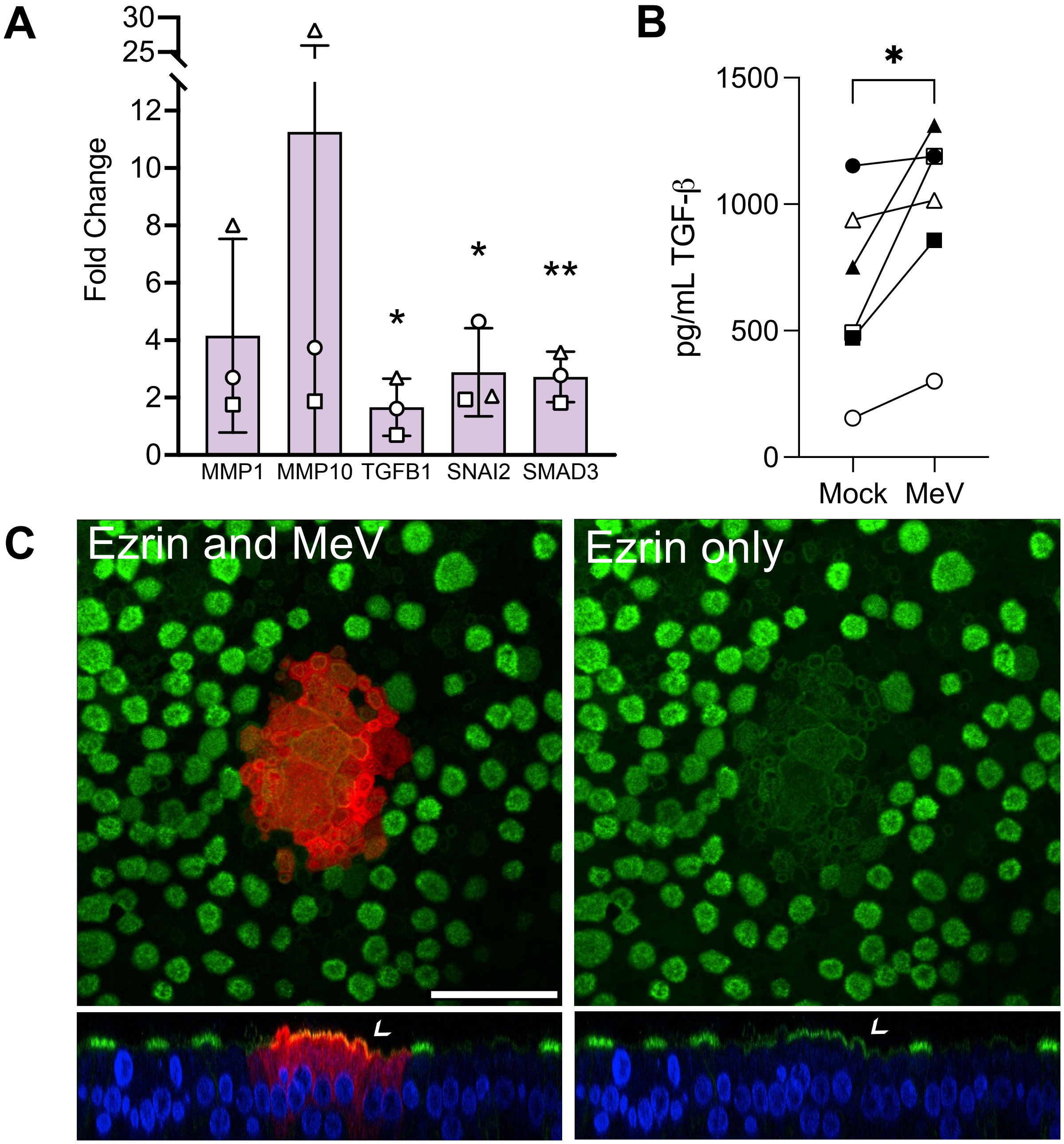

Fig S3. Impact of LCE treatment on infection kinetics.

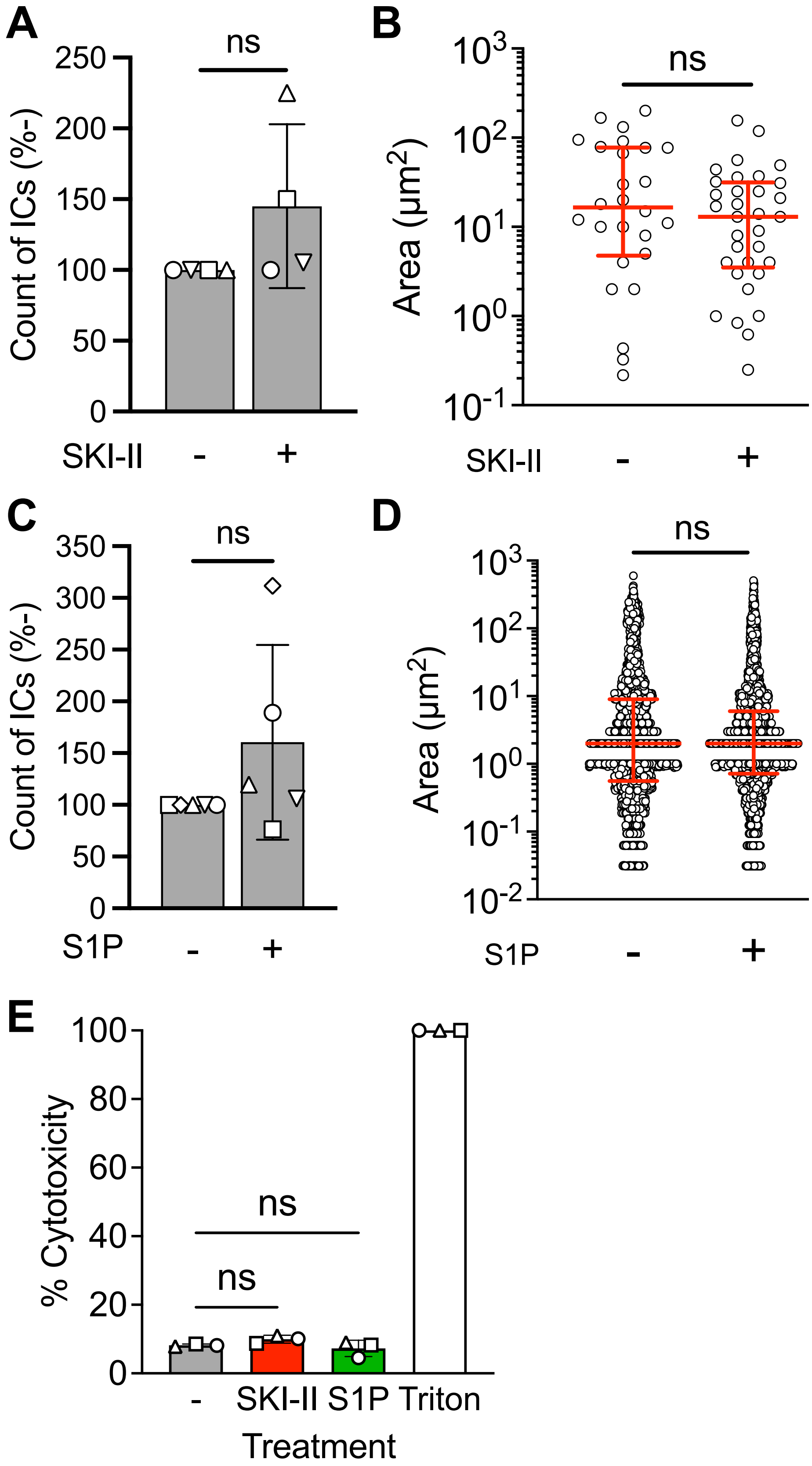
